# *In silico* screen identifies a new *Toxoplasma gondii* mitochondrial ribosomal protein essential for mitochondrial translation

**DOI:** 10.1101/543520

**Authors:** Alice Lacombe, Andrew E. Maclean, Jana Ovciarikova, Julie Tottey, Lilach Sheiner

## Abstract

Apicomplexan parasites cause diseases such as malaria and toxoplasmosis. The apicomplexan mitochondrion shows striking differences from common model organisms, including in fundamental processes such as mitochondrial translation. Despite evidence that mitochondrial translation is essential for parasites survival, it is largely understudied. Progress has been restricted by the absence of functional assays to detect apicomplexan mitochondrial translation, a lack of knowledge of proteins involved in the process and the inability to identify and detect mitoribosomes.

Using mRNA expression patterns, 279 candidate mitochondrial housekeeping components were identified in *Toxoplasma*. 11 were validated, including the mitoribosomal small subunit protein 35 (*Tg*mS35). *Tg*m*S35* tagging enabled the detection of a macromolecular complex corresponding to the mitoribosomal small subunit for the first time in apicomplexans. A new analytical pipeline detected defects in mitochondrial translation upon *Tg*mS35 depletion, while other mitochondrial functions remain unaffected. Our work lays a foundation for the study of apicomplexan mitochondrial translation.

**Abbreviated summary:** The apicomplexan mitochondrion is divergent and essential yet poorly studied. Mitochondrial translation is predicted to utilize ribosomes assembled from fragmented rRNA but this was never shown. Knowing the mitochondrial protein content is critical for these studies. We identified 11 new mitochondrial proteins via *in-silico* searches. Tagging and depletion of a mitoribosomal small subunit protein enabled the first detection of a macromolecular ribosomal complex, and provided proof of principle for our new mitochondrial translation analytic pipeline.

## Introduction

Mitochondria are organelles of central importance to eukaryotic cells, providing key nutrients and metabolites. Mitochondria are present in virtually all eukaryotic cells, excluding a rare case of secondary loss (Karnkowska et al., 2016; Karnkowska and Hampl, 2016). The recent interest in evolutionary cell biology has resulted in an increasing appreciation of the diverse features of mitochondria found in divergent organisms, which includes differences in fundamental pathways such as protein translation (e.g. Ramrath et al., 2018). However, most of our knowledge of mitochondrial biology is still based on studies in organisms that represent a small proportion of the range of eukaryotic diversity, thus limiting our understanding. One of the challenges to the study of mitochondrial biology in diverse organisms is the lack of tools to identify and functionally characterise genes encoding species or phylum specific mitochondrial proteins.

Apicomplexa is a phylum of parasitic protists with high impact on human health globally, which includes the malaria causing *Plasmodium* spp and the causative agent of toxoplasmosis, *Toxoplasma gondii*. Apicomplexans represent one of the lineages whose mitochondrial biology is understudied despite numerous pieces of evidence of its essential role in parasite biology. The apicomplexan mitochondrion hosts an array of essential metabolic pathways (van Dooren et al., 2006; Sheiner et al., 2013). In addition, its central role in parasite survival is highlighted in recent screens for genes important for fitness in *T. gondii* (Sidik et al., 2016) and *Plasmodium berghei* (Bushell et al., 2017). Both screens showed that high numbers of genes encoding mitochondrial proteins are important for fitness. Finally, mitochondrial pathways are the focus of drug discovery studies (e.g. Srivastava et al., 1997; Nilsen et al., 2013; Phillips et al., 2015), including evidence that mitochondrially encoded proteins are promising targets for the development of antimalarials that may circumvent the emergence of drug resistance (Goodman et al., 2016; Klug et al., 2016). It is therefore ever more pressing to expand the studies of mitochondrial biology in these organisms. In this work, we focus on *T. gondii*, a tractable model organism for the phylum.

Some features of the apicomplexan mitochondrion that are divergent compared to the more widely studied yeast, mammal and plant mitochondria have been described. For example, apicomplexans have a single mitochondrion that often spans the entire parasite cell periphery (Seeber et al., 1998; Kudryashev et al., 2010; Ovciarikova et al.,2017). Likewise, the apicomplexan mitochondrion has an extremely reduced genome, with size ranging from 6-11 Kb (Vaidya et al, 1989; Hikosaka et al., 2013), encoding only three proteins. These three proteins are subunits of complexes of the mitochondrial electron transport chain (mETC): cytochrome *b* subunit (Cytb) of complex III, and cytochrome *c* oxidase subunits I and III (CoxI and III) of complex IV (Lin et al., 2011; Feagin, 1992, Suplick et al., 1988, Vaidya et al., 1989, Cinar et al., 2015; He et al. 2014, Ogedengbe et al., 2014). The respiratory chain in apicomplexan differs from more commonly studied mitochondria by lacking complex I (NADH dehydrogenase) and have further different composition of complexes IV (Seidi et al., 2018) and complex V (Salunke et al., 2018; Huet et al., 2018); Finally, mitochondrial translation, which is considered a fundamental pathway in eukaryotes, appears to be highly unusual. For example, *Plasmodium* ribosomal RNAs display a high degree of fragmentation (Feagin et al., 2012; Hillebrand et al., 2018) and it is unclear how these small RNA fragments come together to form a ribosome. Moreover, while evidence for the mitochondrial import of ribosomal and ribosome assembly proteins start to emerge (Ke et al., 2018; Gupta et al., 2018a; Gupta et al; 2018b), no evidence that they form a macromolecular-complex that could correspond to a ribosome has been produced. Nevertheless, it is expected that mitochondrial translation is active in apicomplexans. This was inferred from a combination of findings: a study in *P. falciparum* asexual stages showed that an active mETC is essential for replenishing ubiquinone, which is needed to accept electrons from the parasite dihydroorotate dehydrogenase (DHODH) enzyme (Painter et al., 2007). This was shown by an elegant cross species complementation with the yeast DHODH which uses fumarate as the electron acceptor (Painter et al., 2007). In addition, it was found that resistance to atovaquone, which is an antimalarial drug that inhibits the parasites mETC, is acquired through mutations within the mitochondrial gene encoding the Cytb subunit (Siregar et al., 2008). Together these observations highlight an active mitochondrial translation system essential for the mETC to function and thus for parasite survival. Further support is provided indirectly, by the observed import of tRNAs into the mitochondrion of *T. gondii* and *P. falciparum* (Esseiva et al., 2004, Pino et al., 2010; Sharma and Sharma, 2015). Finally, it was recently shown that a putative mitochondrial ribosomal protein is essential for *P. falciparum* growth, and that its depletion results in mitochondrial biogenesis defects (Ke et al., 2018). However, the observed phenotype upon gene depletion could not be rescued via complementation with yeast DHODH (Ke et al., 2018), leaving some uncertainties about the link between mitochondrial translation and mETC function in *Plasmodium* asexual stages. No direct evidence for this link, or indeed for active mitochondrial translation, is available in *T. gondii*.

Here we report the identification of 11 new *T. gondii* mitochondrial proteins identified in a bioinformatics screen that is based on patterns of mRNA co-expression. Among those we identified an apicomplexan mitochondrial ribosomal protein with homology to small subunit 35, whose function is not studied also in other systems including human. We demonstrate that *Tg*mS35 is part of a macromolecular-complex, providing the first evidence for the assembly of a mitochondrial ribosome in an apicomplexan parasite. We show that *Tg*mS35 is important for parasite growth, and that shortly after *Tg*mS35 depletion a mitochondrial complex, whose subunits are encoded in the mitochondrial genome, shows decreased activity, while other complexes remain active. These observations provide evidence that *T. gondii* mitochondrial translation is active and that it depends on *Tg*mS35.

## Results

### mRNA expression correlation identifies 279 T. gondii genes encoding potential mitochondrial proteins

It was previously shown that patterns of mRNA expression during the *T. gondii* tachyzoite cell cycle have functional predictive power (Behnke et al., 2010; Sheiner et al., 2011). We reasoned that mRNA expression patterns could be used to predict genes encoding components of mitochondrial housekeeping pathways such as translation. At the time of performing this search, the apicomplexan mitochondrial protein import pathway had the most components predicted, we therefore assembled a group of 14 *T. gondii* homologs of these components based on the prediction made in (van Dooren et al., 2006)(Table S1.1, top table). We expected that other parasite mitochondrial housekeeping proteins would share mRNA expression patterns with the genes within this group of search-baits. Accordingly, we queried the microarray data generated from synchronised *T. gondii* tachyzoites with these 14 baits (Behnke et al., 2010). Three genes (*TGME49_274090/260850/251780*) did not show cyclical expression pattern during the cell cycle, therefore we proceeded with the remaining 11 (Table S1.1). We queried the microarray data for mRNAs whose pattern of expression correlates with that of each of these 11 baits. This search identified 279 genes (Figure 1A, Table S1.2, column B).

**Figure 1.**
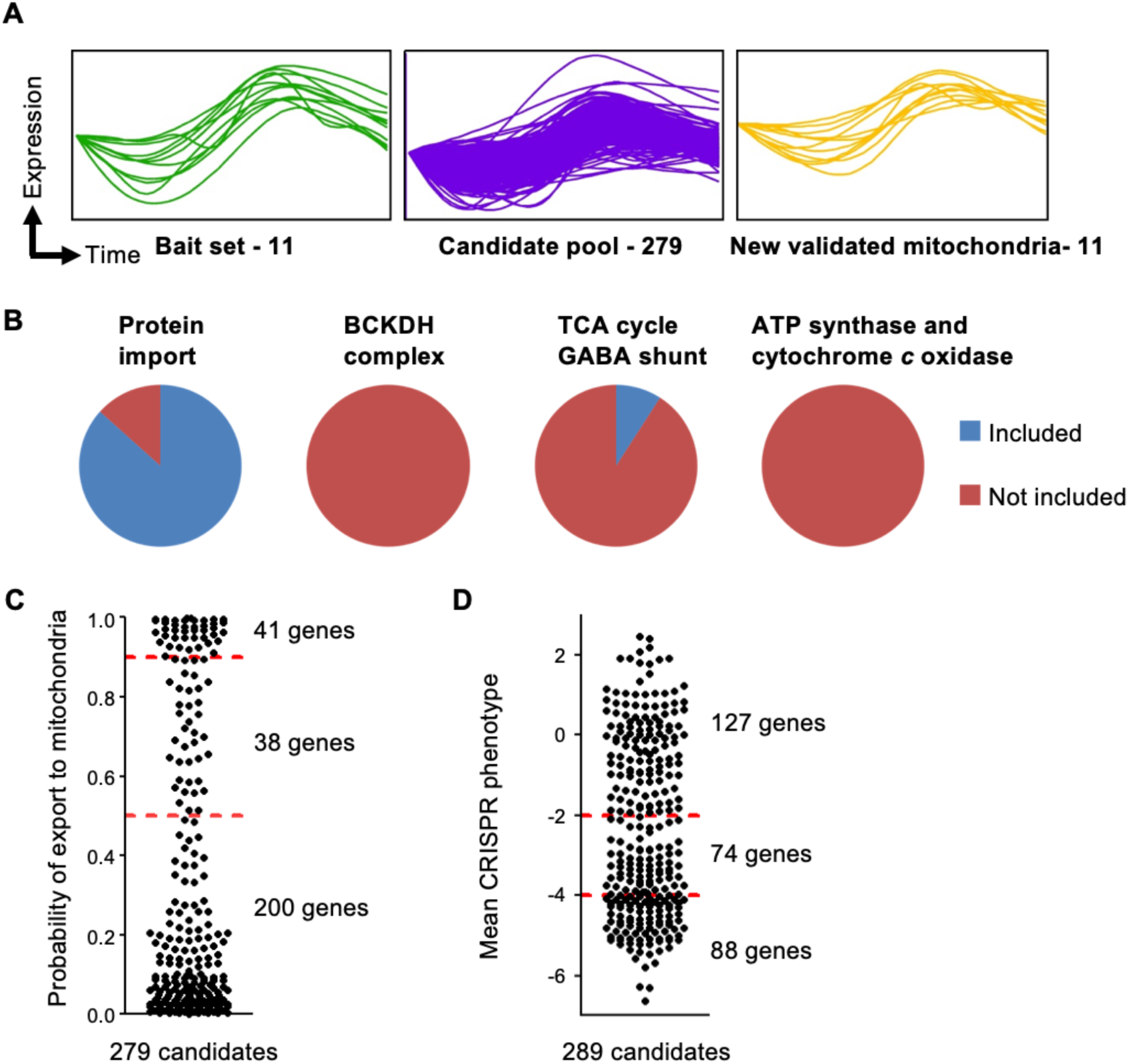
Discovering new mitochondrial proteins using mRNA expression data. (**A**) Graphs show mRNA abundance profiles of the bait set (left, green), candidate pool (middle, purple) and newly validated mitochondrial proteins (right, yellow) over two consecutive tachyzoite cell-cycles. (**B**) Pie charts depicting the inclusion or exclusion of genes from the candidate pool of 279 in functional groups of mitochondrial proteins (protein import, BCKDH complex, TCA cycle and GABA shunt, and ATP synthase and cytochrome *c* oxidase)(geneIDs are in Table S1.3). (**C**) Scatter plot depicting the distribution of the probability of export to the mitochondria of the 279 candidates, assessed using MitoProt II. Candidates with MitoProt score above 0.9 have a high probability of mitochondrial targeting, between 0.5-0.9 a medium probability, and below 0.5 have a low probability. (**D**) Scatter plot depicting the distribution of the mean fitness phenotype scores from the whole genome CRISPR screen of the 279 candidates (genes were first converted from *TGME49* to their orthologous and syntenic *TGGT1* gene models, giving 289 candidates). Genes with a score below −4 are considered “critical”, between −2 and −4 as “important”, and above −2 as “dispensable”. Full datasets for each list are described in Tables S1.2.

Our search criteria were such that each bait was used to query for similar but not identical mRNA expression pattern, namely a search with a given bait does not identify itself. Despite this restriction, the overall resulting dataset includes 10 of the 11 baits used for the searches (TableS1.1, column E), meaning that queries with some baits identified the other baits, providing validation for this approach. Moreover, another four genes encoding protein import components that were not used in the bait list (since they were not predicted at the time the search was performed) are also found in this dataset (TableS1.1, bottom table). Additionally, genes encoding mitochondrial proteins that function in other known (metabolic and physiological) pathways that are not expected to co-express with the mitochondrial protein import encoding genes are indeed underrepresented in the resulting dataset (Figure 1B, TableS1.3, columns I-R), validating this approach’s specificity. We observed that 28.3% are predicted to poses a mitochondrial targeting signal with probability score of >0.5 (Figure 1C). We further observed that 56% of the identified genes are predicted to be important for fitness (scores >-6.8 and <-2 in the whole genome CRISPR screen (Sidik et al., 2016)) (Figure 1D, Table S1.3, Column E) compared to 40% predicted to contribute to parasite fitness in the whole genome screen study (Sidik et al., 2016), or 24.824% (2,875 of total 8,637 genes) if running the above score criteria in ToxoDB. Enrichment of genes important for fitness is in line with enrichment in mitochondrial genes, given that they are highly represented in fitness screens (Sidik et al., 2016; Bushell et al., 2017). In contrast, only 39% of the 279 are unique to apicomplexan (Table S1.3, column C) compared to 42.3% (4,899 of total 8,637 genes) in the whole genome, which is only a minor difference.

### Among 15 genes selected from the 279-gene dataset, 11 encode mitochondrial proteins

As a proof of principle, we chose 10 genes encoding hypothetical proteins to assess their localisation (Table 1). We primarily used endogenous gene tagging via single homologous recombination to introduce an epitope tag fusion at the protein c-terminal end (Huynh and Carruthers, 2009; Sheiner et al., 2011). Where endogenous tagging was unsuccessful we assessed the localisation via expression of the tagged cDNA from a heterologous promoter. Among the 8 genes for which we obtained detectable signal (Table 1), 7 were mitochondrial (Figure 2A).

**Table 1.** Localisation and sequence-based analysis of the 15 genes experimentally studied from the 279 candidates. Phylogenetic observations are based on search in OrthoMCL via ToxoDB and on tBLASTn via NCBI. ^OrthoMCL and BLAST search do not identify homologs outside apicomplexans or alveolates, however evidence is available for homologs outside these groups. Empty cells are where homologs are found outside apicomplexan. NA – we did not look at phylogenetic distribution when the gene product was not localised to the mitochondrion. Mitochondrial targeting predictions are done using MitoProt II (https://ihg.gsf.de/ihg/mitoprot.html) and Predotar (https://urgi.versailles.inra.fr/predotar/). The presence (yes) or absence (no) of the gene product in the proximity tagging based mitochondrial proteome is shown (Seidi et al., 2018) (no - no mass spectrometry data; no* - mass spectrometry data found but the gene is not included in the final list of 421 genes). The fitness score is based on the whole genome CRISPR/CAS9 screen (Sidik et al., 2016) (geneIDs in bold and with ^§^-essentiality/importance for fitness confirmed here).

| Gene ID | Predicted product (TGME49) | Localisation Form of tagging | Phylogeny comment | MitoProt | Predotar | Proximity tagging proteome | Phenotype score |
| --- | --- | --- | --- | --- | --- | --- | --- |
| **10 genes with hypothetical gene product from the 279-gene dataset** |  |  |  |  |  |  |  |
| **TGME49_203620<sup>s</sup>** | hypothetical protein | Mitochondrial cDNA | Apicomplexan unique<sup>^</sup> | 0.9734 | 0.53 | no* | -4.66 |
| TGME49_226270 | hypothetical protein | Apicoplast endogenous | NA | 0.0402 | 0.01 | no | 0.23 |
| TGME49_231150 | hypothetical protein | Mitochondrial endogenous |  | 0.9986 | 0.33 | yes | -5.21 |
| **TGME49_240270<sup>s</sup>** | hypothetical protein | Mitochondrial cDNA | Apicomplexan unique | 0.2831 | 0.04 | no* | -4.65 |
| TGME49_240780 | hypothetical protein | Mitochondrial endogenous | Apicomplexan unique | 0.3871 | 0.76 | no | -3.95 |
| TGME49_247740 | hypothetical protein | Mitochondrial endogenous | apicomplexan and bacteria | 0.0647 | 0.03 | no* | 0.31 |
| TGME49_257110 | hypothetical protein | no signal cDNA | NA | 0.6373 | 0.01 | no | -3.21 |
| TGME49_263680 | hypothetical protein | Mitochondrial endogenous | Apicomplexan unique | 0.9490 | 0.52 | no* | -4.30 |
| TGME49_270150 | hypothetical protein | Mitochondrial endogenous | Apicomplexan unique | 0.9859 | 0.67 | yes | -4.23 |
| TGME49_310500 | hypothetical protein | no signal cDNA | NA | 0.9211 | 0.11 | no* | -4.10 |
| **5 genes with chosen from the 43-gene dataset** |  |  |  |  |  |  |  |
| TGME49_201830 | ribosomal protein L37 | Mitochondrial endogenous | Alveolata unique<sup>^</sup> | 0.9991 | 0.43 | no | -4.94 |
| TGME49_226280 | ribosomal protein L28, putative | Mitochondrial endogenous |  | 0.9971 | 0.65 | yes | -4.91 |
| **TGME49_214790<sup>s</sup>** | glycoprotein | Mitochondrial endogenous |  | 0.8383 | 0.47 | yes | -3.48 |
| TGME49_311750 | mago binding protein | Cytosol cDNA | NA | 0.1980 | 0.01 | no | -0.99 |
| TGME49_312680 | 60S ribosomal protein L27, putative | Mitochondrial cDNA |  | 0.9388 | 0.14 | no | -1.34 |

**Figure 2.**
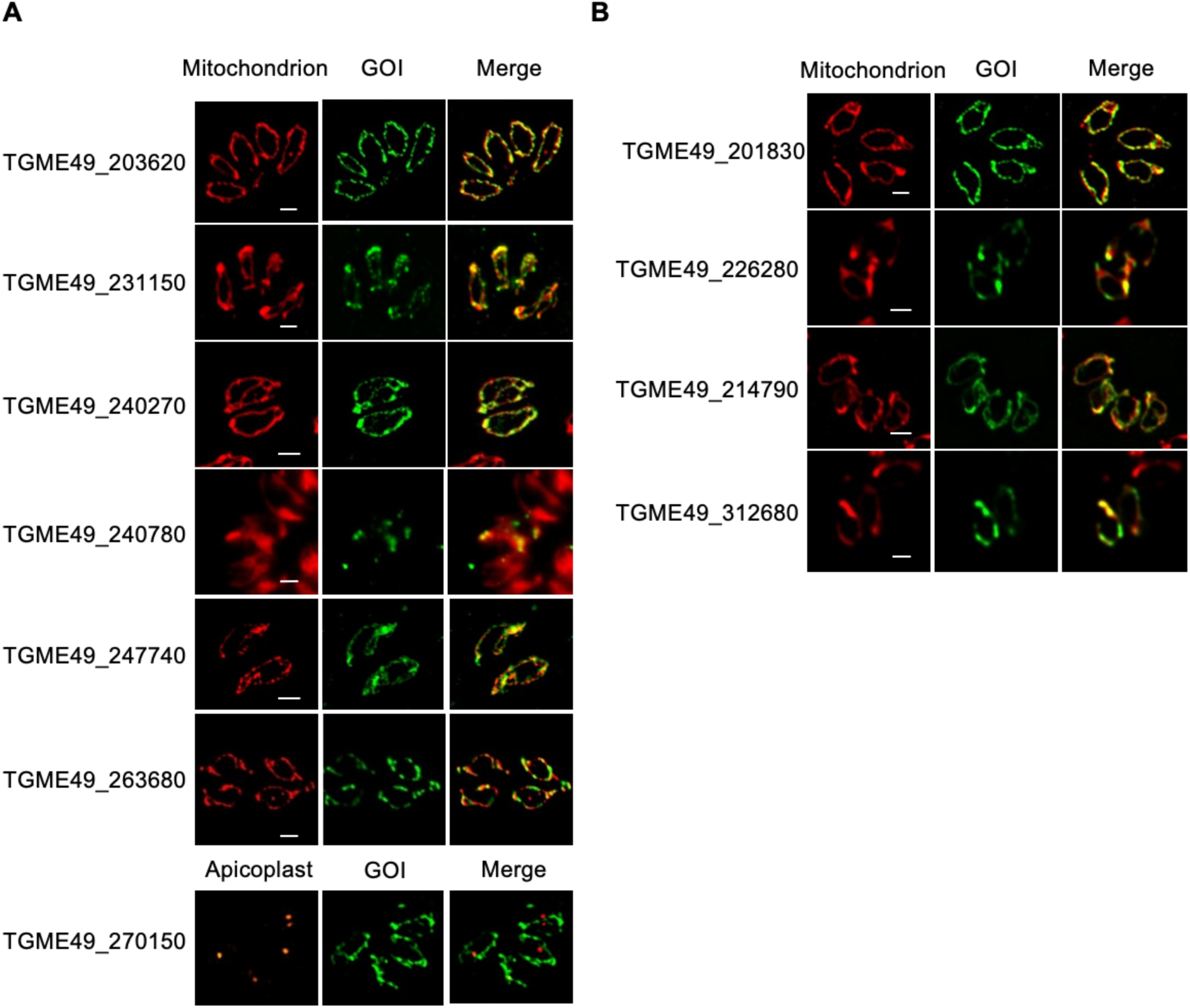
Mitochondrial localisation of 11 previously uncharacterised proteins by epitope tagging. (**A**) Localisation through fluorescence microscopy analysis of 7 gene-products annotated as “hypothetical proteins” from the candidate list of 279 genes. (**B**) Localisation of 4 gene-products from the candidate list of 43 genes with no homologs in *Cryptosporidium* spp. For both panels, mitochondria are marked with anti-TgMys or anti-TOM40; the mentioned GOI (gene of interest) are marked with HA or YFP; apicoplast is marked with anti-CPN60; Scale bar 1 µm.

To sharpen our focus on understudied mitochondrial housekeeping pathways such as mitochondrial translation, we added another filter to our search: we postulated that *Cryptosporidium* spp, that are the only known species among the apicomplexans that lost their mitochondrial genome, likely lost their mitochondrial translation pathway, that is otherwise conserved among other apicomplexans. We queried the datasets for genes that do not have orthologs in any of the three genomes available for *Cryptosporidium* spp, but that do have orthologs in the genome of *Plasmodium falciparum* 3D7. This search identified 43 genes (Table S1.2, column C). In support of an enrichment of translation components in this dataset, we found 5 of the 35 predicted mitochondrial ribosomal proteins (Gupta et al., 2014) in the 43-gene dataset, while none of them is found in 5 lists of 43 randomly chosen genes (Table S1.3, column U). We attempted to localise five genes from this focused dataset chosen at random, and four localised to the mitochondrion (Figure 2B).

### Transient expression of CAS9 results in mitochondrial morphology defects

Having a group of 11 new mitochondrial proteins to study, we wanted to examine whether transiently expressed CRISPR/Cas9 strategy could be used to provide a rapid read-out assay to detect protein’s involvement in mitochondrial biogenesis. We hypothesised that abnormal changes in mitochondrial morphology upon gene disruption may be an indicator for a defect in mitochondrial biogenesis. We picked three of the new mitochondrial proteins at random (encoded by *TGME49_263680/214790* and *226280*). For each gene, we co-expressed FLAG-tagged Cas9 (Sidik et al., 2014) along with a targeting single guide RNA (sgRNA)(TableS2). As control, we expressed the Cas9-FLAG alone or with sgRNA for the non-essential SAG1 (*TGME49_233460*) gene for which we do not expect any mitochondrial phenotype upon gene disruption (SAG1sgRNA). At 48 hours post co-transfection, mitochondria were imaged using immunofluorescence and morphologies were analysed (Figure 3A). We observed elevated mitochondrial abnormalities in all conditions tested except the untreated parasites. While a trend of higher abundance of mitochondrial abnormalities was observed for *TGME49_214790*, the data showed no significant difference from the Cas9-only and Cas9+SAG1sgRNA controls (Figure 3A, B). These data suggest that transient expression of Cas9 results in mitochondrial morphological abnormalities, excluding this strategy as a reliable tool for rapid identification of genes involved in mitochondrial biogenesis.

**Figure 3.**
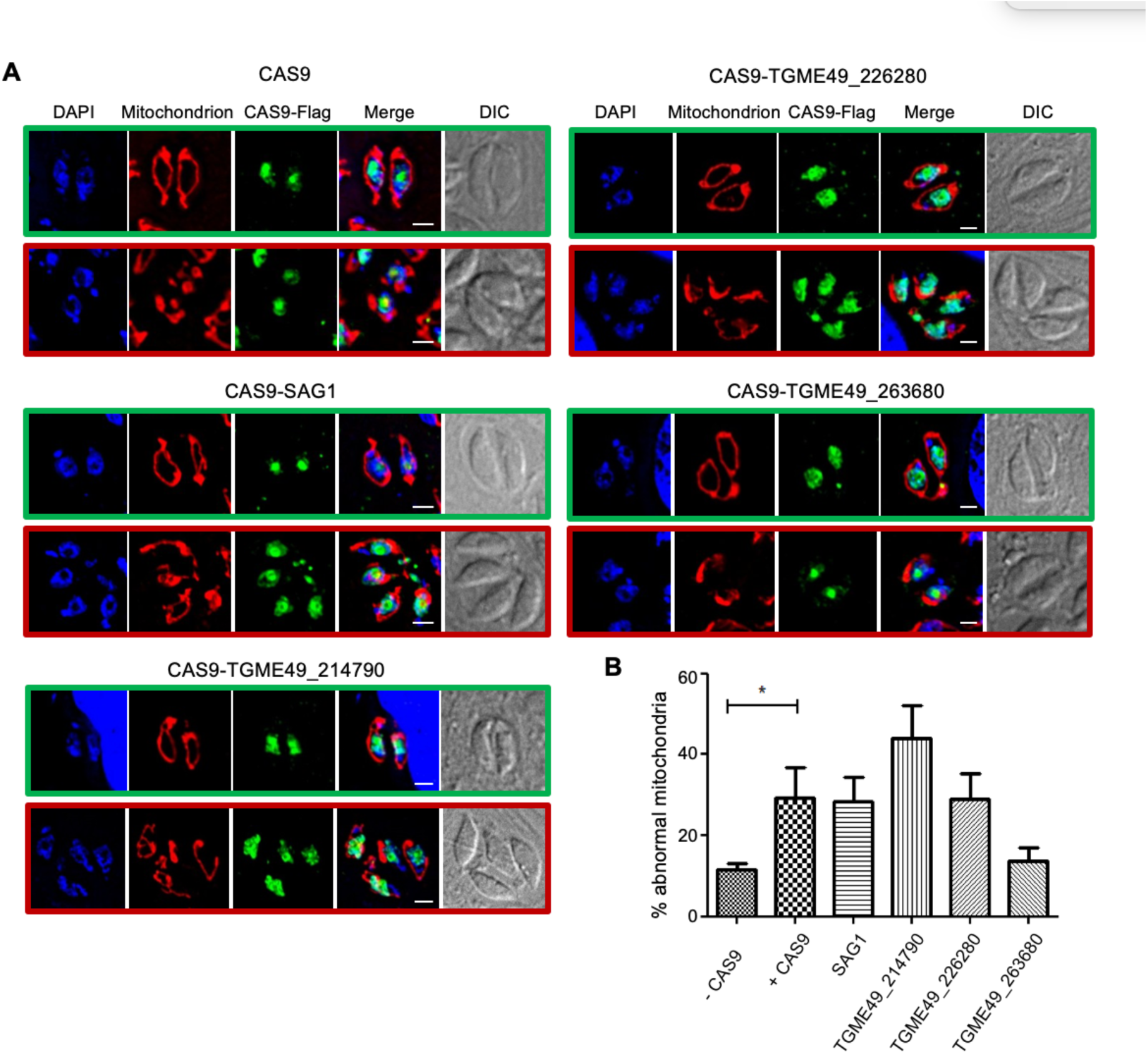
Transient expression of Cas9 results in mitochondria morphology defects. (**A**) Representative immunofluorescence micrographs of parasites transiently expressing CAS9 and sgRNA for the mentioned GOI. Mitochondria in red are marked by anti-TgMys. CAS9-FLAG in green is marked with anti-FLAG. Merge shows DAPI, FLAG and TgMys. Green boxes highlight parasites with wild-type looking mitochondria. Red boxes highlight parasites with mitochondria that appear to have a morphological defect herein named “abnormal mitochondria”. (**B**) Quantification of vacuole with abnormal mitochondria morphologies for each condition. Bars represent the mean ± SEM (n=3), * p < 0.05 (1-way ANOVA).

### *TGME49_203620* encodes a ribosomal protein crucial for parasite fitness

Due to the complication encountered in analysing mitochondrial biogenesis defects in parasites transiently expressing Cas9, we proceeded with functional analysis using stable genetic manipulation. As a means to explore the role of mitochondrial translation in *T. gondii* survival, we chose to focus on a gene encoding a ribosomal protein. One of the genes found in our screen, *TGME49_203620*, while annotated as hypothetical protein in ToxoDB.org, is a putative homolog of the mitochondrial small subunit 35 (mS35) (Gupta et al., 2014; Greber and Ban, 2016). To provide support for this prediction we tested the gel migration patterns of endogenously triple-HA tagged *TGME49_203620*. While *TGME49_203620*’s protein product migrates around 38 KDa under denaturing conditions (Figure 4A), around the predicted size of the part of the protein that is conserved among apicomplexans (Figure S1), it migrates with a high molecular weight (>1000 KDa) complex, when separated under native conditions using blue-native PAGE (Figure 4B). The migration under native conditions could well correspond to the mitochondrial ribosomal small subunit complex. We therefore named *TGME49_203620*, *Tg*m*S35*.

**Figure 4.**
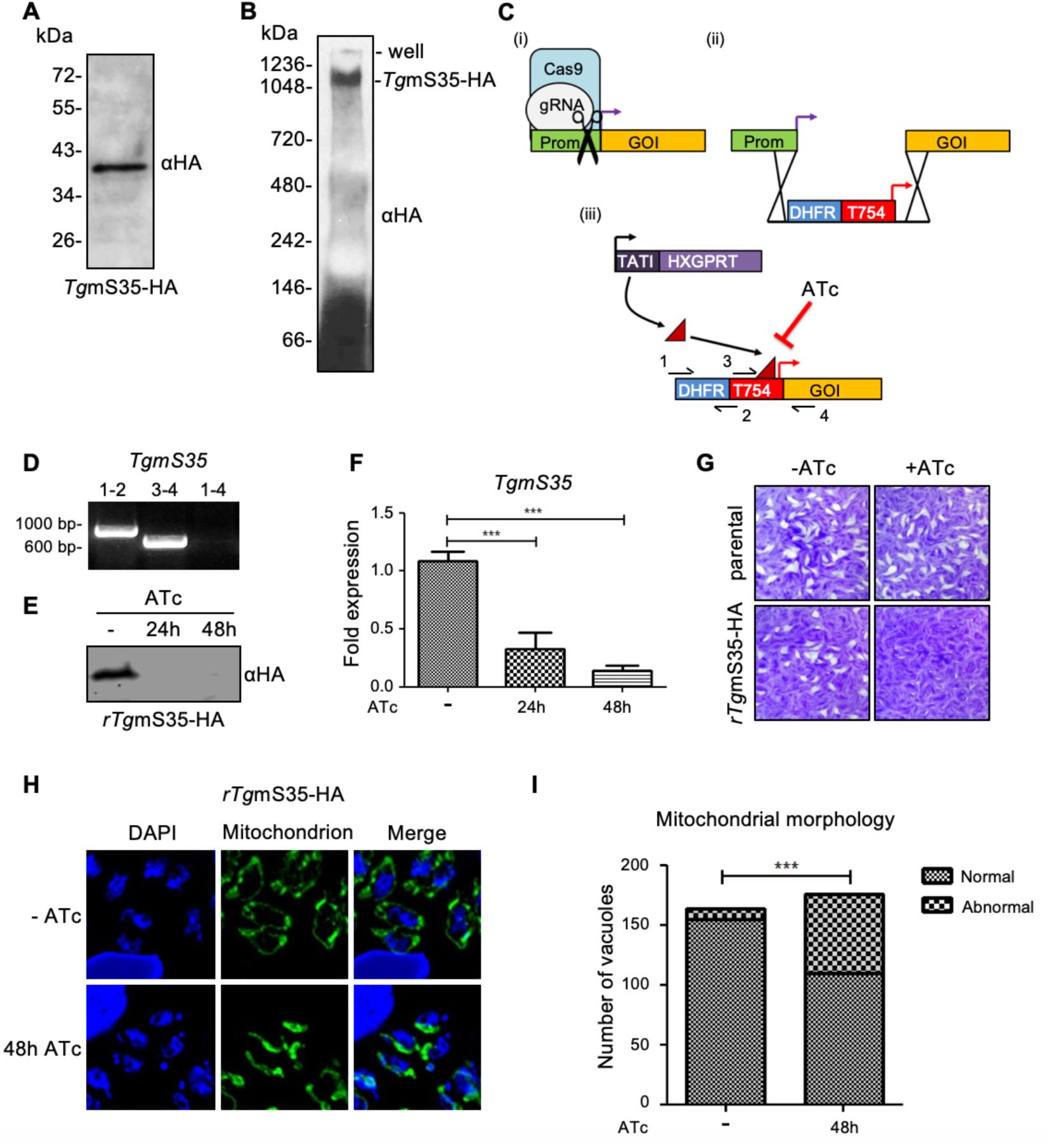
Knock-down and analysis of *TGME49_203620*, a homolog of the mitoribosomal subunit *mS35*. (**A**) Protein immunoblot analysis of endogenously 3xHA tagged *Tg*m*S35* (TGME49_203620) from total cell lysate separated by SDS-PAGE and detected using an anti-HA antibody. (**B**) Total cell lysate separated by blue-native PAGE and immunoblotted to detect *Tg*mS35-3xHA with an HA antibody. (**C**) Schematic depiction of the promoter replacement strategy allowing knock-down of a GOI with the addition of ATc. (i) CRISPR/CAS9 guided cut at the predicted promoter/ATG boundary, (ii) a repair cassette containing the ATc repressible promoter, the DHFR selection marker, and homology to the promoter/ATG boundary, is inserted between the promoter and GOI, (iii) GOI, under the control of ATc repressible promoter, is down regulated when ATc is added. The black arrows represent the four primers used to confirm integration. (**D**) Validation of the promoter integration in the *Tg*mS35 locus via PCR analysis using primers 1, 2, 3, and 4, represented in C (iii). (**E**) Protein immunoblot analysis of whole cell lysate from r*Tg*mS35-HA after growth in absence (-) or presence of ATc for 24 and 48 hours. (**F**) Transcript levels of *Tg*mS35, analysed by qRT-PCR, in absence (-) or presence of ATc after 24 and 48 hours. Bars represent the mean ± SEM (n=5). ***p< 0.0001 (**G**) Plaque assays performed with parental line or r*Tg*mS35-3xHA parasites grown for 9 days in the absence (-) or presence (+) of ATc. (**H**) Immunofluorescence micrographs taken in the presence or absence (-) of ATc for 48h, showing examples for wild type (-ATc) or abnormal (48h ATc) mitochondrial morphologies. (**I**) Quantification of normal vs abnormal mitochondria containing vacuoles after growth of r*Tg*mS35 without (-) or with ATc for 48 hours (n=3); ***p< 0.0001 (Chi-squared test).

We generated a *Tg*m*S35* conditional knock down line by replacing its native promoter with an anhydrotetracycline (ATc) repressible promoter. This manipulation was performed as previously described (Sheiner et al., 2011), however with the modification whereby CRISPR/Cas9 was utilised to enhance integration of the promoter replacement cassette (scheme Figure 4C). We named this line *rTgmS35-3xHA*. Inducible promoter integration was confirmed by PCR (Figure 4D) and ATc induced down-regulation was confirmed by qPCR and by western blot using the triple HA tag (Figure 4E-F). We monitored parasite growth by plaque assay which revealed a severe growth defect in response to *Tg*m*S35* depletion, however small plaques revealed that parasites can still grow slowly (Figure 4G). Finally, immunofluorescence analysis detected mitochondrial morphological abnormalities upon *Tg*m*S35* depletion (Figure 4H-I). Together, these results are consistent with *Tg*m*S35* being a mitochondrial ribosomal subunit important for parasite fitness and mitochondrial biogenesis. Using the same genetic manipulation strategy, we generated conditional mutants for two additional genes among the 11 newly identified mitochondrial protein encoding genes and showed that both are important for parasite growth (Figure S2).

### A new protocol for evaluation of mitochondrial translation shows that *Tg*mS35 is essential for this pathway

In yeast, mS35 is found at the critical point of mRNA entry into the ribosome (Desai et al., 2017). We hypothesised that depletion of *Tg*m*S35* should result in a mitochondrial translation defect, however the field lacks an assay to directly monitor mitochondrial translation in *T. gondii*. We therefore monitored the assembly and activity of the respiratory chain complex IV, which is directly dependent on the mitochondrial translation of CoxI and III, and compared it to the assembly and activity of complex V, which is not dependent on mitochondrial translation of its subunits. Using an enriched mitochondrial fraction, obtained from parasites lysed through nitrogen cavitations, we performed high resolution clear-native PAGE followed by an in-gel complex IV and V enzymatic assays, at different time points after *Tg*m*S35* depletion (Figure 5A, B). We found a decrease of complex IV activity at 48 hours and loss of the activity at 96 hours after *Tg*m*S35* depletion. In contrast, the activity of complex V is unchanged at 48 hours and is only decreased at the 96-hour time point indicating an indirect defect (Figure 5A, B). The band observed for complex IV enzymatic activity migrates at a size consistent with a previous report (Seidi et al., 2018). To further confirm its identity as complex IV we performed mass spectrometry analysis of this band which confirmed the complex identity (Table S3).

**Figure 5.**
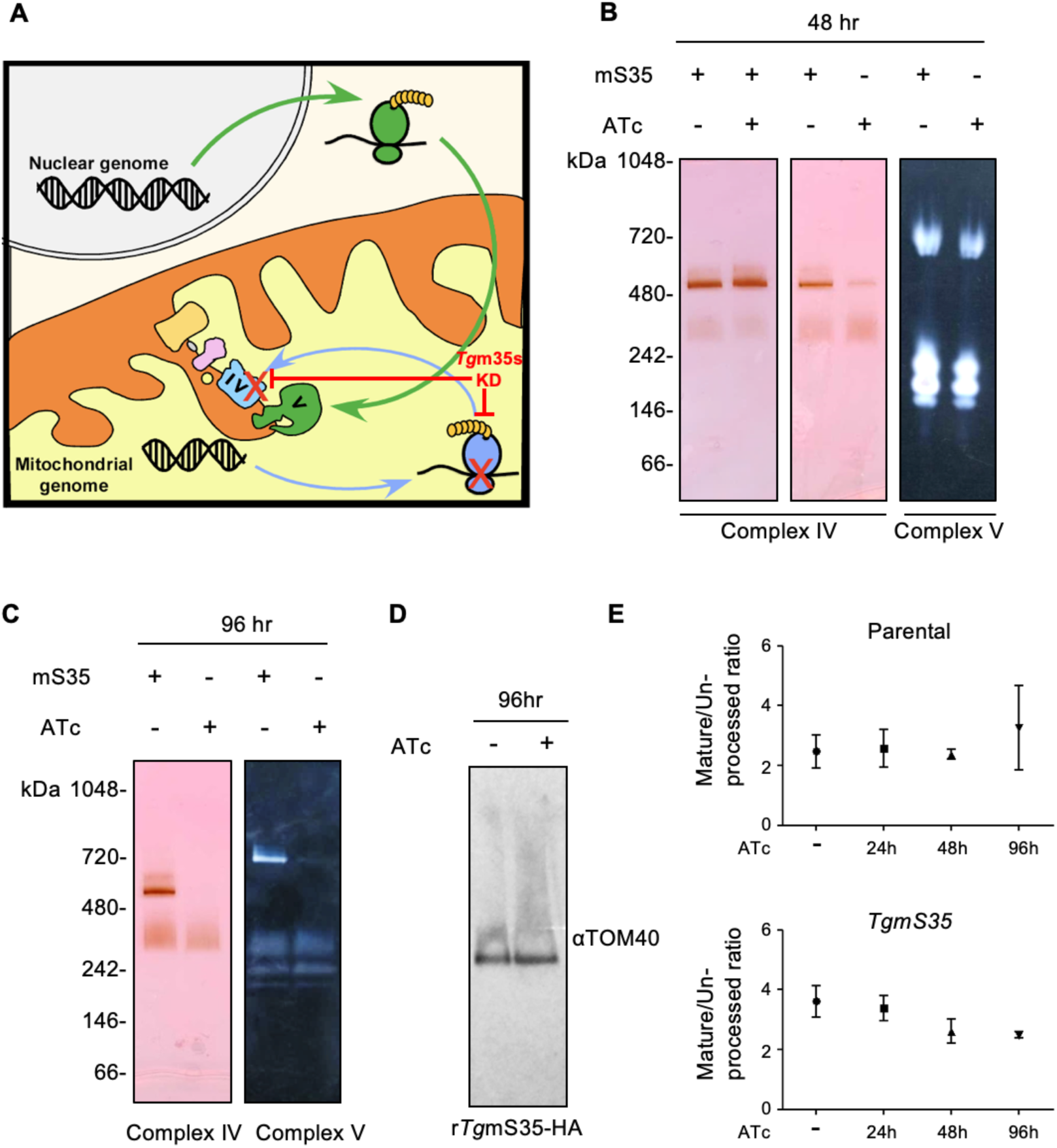
Down-regulation of *Tg*mS35 leads to a defect in respiratory complex IV. (**A**) A scheme describing the rational of our assay. Nuclear (grey) encoded proteins translated by the cytosolic ribosome (green) compose complex V, while mitochondrial (yellow) encoded proteins translated by the mitochondrial ribosome (blue) are necessary to assemble complex IV. Depletion of Tgm35S (red) results in reduced activity of complex IV but not V. (**B**) Whole cell (left panel) and enriched mitochondria (middle and right panel) from *Tg*mS35-3xHA (left panel) and from r*Tg*mS35-3xHA (middle and right panel), grown in the absence (-) or presence (+) of ATc for 48 hours, separated by high resolution clear-native PAGE. Complex IV activity was visualised with cytochrome *c*: DAB staining. Complex V activity was assayed by visualising the precipitation of lead by inorganic phosphate during ATP hydrolysis. Protein immunoblot analysis using anti-TOM40 of whole-cell and enriched mitochondria sample was performed as a loading control. (**C**) Mitochondrial enriched fraction from r*Tg*mS35-3xHA grown in the absence (-) or presence (+) of ATc for 96 hours, separated by high resolution clear-native PAGE and stained with coomassie or for complex IV or complex V activity as above. **(D)** Immunoblot with anti-TOM40 antibody of mitochondrial enriched fraction from r*Tg*mS35-3xHA grown in the absence (-) or presence (+) of ATc for 96 hours, separated by blue-native PAGE. **(E)** Quantification of the ratio of bands of un-processed and mature version of Hsp60_L_-mDHFR-cMyc measured in an import assay in parental (top graph) and r*Tg*mS35-3xHA (bottom graph) parasites lines in the absence (-) or presence of ATc for 24, 48 and 96 hours. Bars represent the mean ± SEM (n=3 for the parental line, and n=5 for r*Tg*mS35-3xHA).

In contrast to complex IV, western blot analysis of a native gel from the 96-hour time point showed that the protein import complex, translocon of the outer membrane (TOM), migrates at the same size as the untreated line (Figure 5C). This data suggests that TOM complex, whose components are not encoded in the mitochondrial genome, continues to assemble. To provide further support for TOM complex assembly, we measured protein import into the mitochondrion. A Myc tagged DHFR fused to the mitochondrial targeting signal of HSP60 (Hsp60_L_-mDHFR-cMyc (van Dooren et al., 2016)) was transiently transfected into the parasites at different time points after *Tg*m*S35* depletion. Upon entry into the mitochondrion, the mitochondrial targeting signal is cleaved off, generating a smaller sized protein which can be visualised by western blot. Upon an import defect, cleavage no longer takes place (van Dooren et al., 2016). We calculated the ratio of signal intensities between the cytosolic un-cleaved protein and the mitochondrial cleaved protein in each condition. We did not observe any significant protein import defect upon *Tg*m*S35* depletion, including at the 96-hour time point (Figure 5D). Taken together, these observations suggest that *Tg*m*S35* depletion results in a defect in the assembly and activity specifically of a mitochondrial respiratory complex whose subunits are thought to be encoded in the mitochondrial genome.

## Discussion

Research in recent years has expanded our understanding of mitochondrial diversity across the eukaryotes. However, there is still a reliance on a handful of models, such as yeast and mammals or such as plants, when studying basic mitochondrial functions. Focusing on these canonical mitochondria gives a skewed view of mitochondrial function and composition. One of the reasons for this focus is the lack of well-developed, tractable, model organisms outside these key groups. Despite key advances in recent years in studying microbial eukaryotes, there is still a lack of datasets and tools for studying mitochondrial biology in non-model groups. Apicomplexa, which contain the medically important *Plasmodium* and *Toxoplasma* parasites, is such a group where the mitochondria are understudied. The herein described work provides crucial advances by generating proof of principal for the functional prediction of mRNA expression patterns and by establishing an analytical pipeline to identify mitochondrial translation defects.

### Co-expression patterns predict new mitochondrial proteins while also providing functional insight about the resulting candidates

To fully understand and explore mitochondrial functions it is necessary to create an inventory of mitochondrial proteins. To date only 88 *Toxoplasma* mitochondrial proteins have been experimentally validated (summarised in (Mallo et al., 2018)). Establishing the proteome through organelle isolation has been a challenge to the apicomplexan field partially due to the association between the mitochondrion and the apicoplast (Kobayashi et al., 2007; Nishi et al., 2008; and our unpublished data). While methods that may resolve this complication are being explored (Hata et al., 2018), other strategies that bypass the reliance on organelle isolation are being utilised. Recent biotin-proximity tagging approaches have made a big step forward (Seidi et al., 2018). However, the resulting dataset covers mainly matrix proteins. Moreover, the number of 421 proposed mitochondrial proteins identified is predicted to be markedly smaller than the total number of mitochondrial proteins estimated through comparison to other species (Salvato et al., 2014; Smith and Robinson, 2018; Panigrahi et al., 2009). Here we demonstrated the power of mRNA expression pattern in *T. gondii* to identify novel mitochondrial components, and thus to expand the repertoire of known mitochondrial proteins. Moreover, a limitation of strategies aimed to create inventories, is that the resulting datasets do not provide functional insights. We provide evidence that the expression-based prediction provides functional insights, as our chosen group of baits enriched for proteins involved in mitochondrial housekeeping. In this, our finding joins previous work in showing that mRNA expression patterns in *T. gondii* are powerful predictive tool for genes in shared functional groups. Behnke et al, who generated the mRNA expression data used for this study, showed that genes known to be involved in the same pathways (e.g. nuclear genome maintenance, microneme adhesins, rhoptry effectors) share mutual expression pattern (Behnke et al., 2010). Likewise, this strategy was used to predict proteins involved in host cell invasion (Huynh and Carruthers, 2016). Moreover, the opposite is also true, when the group of baits is not composed of proteins that take part in the same functional pathway, the resulting dataset contains proteins from different functional pathways as we showed for apicoplast proteins (Sheiner et al., 2011). Only 4 of the 11 new mitochondrial proteins identified here are predicted in the Seidi et al., dataset of 421 genes (Table 1). It is possible that the dataset generated here has higher representation of proteins from non-matrix mitochondrial sub compartments (A potential example is TGME49_240270, which is not predicted to have a mitochondrial targeting signal (Table 1)), or of genes encoding non-abundant proteins that may be missed by proteomics analyses (e.g. TGME49_240780, TGME49_201830 and TGME49_312680 (Table 1)).

Our study focused on mRNAs that co-express with homologs of the mitochondrial protein import machinery components, some of which were recently validated experimentally (van Dooren et al., 2016). In the same study the authors identified new components of these import complexes that were not included in our bait list (van Dooren et al., 2016). The presence of three of these five new import components in our 279-gene dataset (Table S1.1, bottom table) lends support to the power of this strategy to identify mitochondrial protein import components, especially considering none of them are found in five lists of randomly chosen 279 genes (Table S1.4, columns B-M). This observation raises the possibility that some other genes in the dataset may take part in protein import. A previous study showed that at least one of the mitochondrial protein translocation complexes has comparable size to other organisms (van Dooren et al., 2016), while homologs of many components known in other organisms are missing in *T. gondii* (van Dooren et al., 2016), suggesting that currently unknown components are part of this machinery and that they may be parasite specific. The 109 apicomplexan unique genes in our dataset (Table S1.3, column C) provide candidates for those missing pieces. Another pathway whose components may co-express with protein import components is mitochondrial tRNA import (Esseiva et al., 2004; Pino et al., 2010; Sharma and Sharma, 2015), as in some organisms these two pathways are linked (Seidman et al., 2012; Tschopp et al 2011). Mitochondrial tRNA import is likely essential for mitochondrial translation, for which components are enriched in the 43-gene dataset, proposing these 43 genes as candidate for component of this pathway.

Localisation of proteins encoded by 13 candidates from the 279-gene dataset, for which a signal was detected by immunofluorescence, identified 11 new *T. gondii* mitochondrial proteins, expanding the group of validated mitochondrial proteins by 11%. Sequence analysis of the 11 proteins demonstrate that our search strategy provides some improvement from the available mitochondrial targeting prediction algorithms, as either two or three of the new mitochondrial proteins are not predicted to target the mitochondrion in two different algorithms (Table 1). These finding highlight the limitation of relying on localisation prediction algorithms for the prediction of apicomplexan mitochondrial proteins. A similar observation was reported in the biotin proximity study (Seidi et al., 2018). This may be one of the reasons for the observation that more than 70% of the 279 proteins have low score (under 0.5) for the prediction of mitochondrial targeting when using the MitoProt II algorithm (https://ihg.gsf.de/ihg/mitoprot.html) (Claros and Vincens., 1996). Another reason for this observation is that many of the non-matrix proteins in the dataset utilise other targeting signals. For example, any mitochondrial proteins among the 50 in the dataset that have predicted transmembrane domains (Table S1.3, column F) likely enters the mitochondrion independently of a mitochondrial targeting signal.

### Mitochondrial morphology captured by immunofluorescence is linked to mitochondrial functional defects in *Toxoplasma*

In model organisms, mitochondrial morphology and function are tightly linked. In support of this also being the case in *Toxoplasma*, recent studies reported rapid mitochondrial remodelling in respond to drastic changes in the growth environments (Ovciarikova et al., 2017; Charvat and Arrizabalaga, 2016). Likewise, the morphological defect observed here upon *Tg*mS35 depletion provides additional support. For this reason, we hoped to use mitochondrial morphological changes as a preliminary indicator for mitochondrial functional defects. However, we found that transient expression of the Cas9 endonuclease in *T. gondii* induces mitochondrial morphology defects. This observation should be considered when analysing mitochondrial phenotypes under these conditions.

### Observation of the mitochondrial ribosome and characterisation of a new ribosomal component demonstrate the crucial role of mitochondrial translation in *T. gondii*

A recent structure of the mitochondrial ribosome of *Trypanosoma brucei* showed an evolutionary shift towards a higher ribosome protein content rather than the RNA heavy complexes of mammalian ribosomes (Ramrath et al., 2018). These findings from *T. brucei* illustrates the importance of studying basic cellular components in a diverse array of models, however, this is not possible without detecting mitochondrial ribosomes and validating mitochondrial ribosomal proteins. Among the 11 genes we localised here, 4 are predicted to be ribosomal subunits. One of those, *TGME49_203620*, while annotated in some databases as hypothetical, shows homology to the mitochondrial small ribosomal subunit 35, mS35, and was therefore renamed *Tg*mS35. Blue-native separation of endogenously tagged *Tg*m*S35* resolved it as a part of a macro-molecular complex, with a size potentially corresponding to the small mitochondrial ribosomal subunit complex. This is the first direct evidence that, despite the apicomplexan fragmented mitochondrial rRNA (Barbrok et al., 2010), the apicomplexan mitochondrial ribosomes are assembled into stable complexes. This is a major step forward in the study of apicomplexan ribosomes which are not well studied and whose composition is not known. While our observation suggests that macro-molecules containing mitochondrial ribosomal proteins form a stable complex, the question of how they assemble and are held together structurally remains open. A recent study identified novel RNA binding proteins in the apicomplexan mitochondrion and proposed the hypothesis that they may participate in ribosome assembly (Hillibrand et al., 2018). The tagged ribosomes we generated here can be used in the future to try tackling the question of mitochondrial ribosome structure and function in *T. gondii*.

### A new protocol to follow mitochondrial translation in *T. gondii*

The lack of a direct assay to detect mitochondrial translation in Apicomplexa is a major obstacle to study this important pathway. In a recent study, a *P. falciparum* mitochondrial ribosomal protein, the large subunit 13 (which per the new notation (Greber and Ban., 2016) can be named PfuL13m) was characterised (Ke et al., 2018). Like *Tg*mS35, the study found that PfuL13m depletion mutant displayed a growth defect. The authors showed decreased activity of respiratory complex III, whose Cytb subunit is mitochondrially encoded, as a read-out for translation (Ke et al., 2018). However, the inability to rescue the growth defect using an mETC bypass route available for the *Plasmodium* asexual stages (Ke et al., 2018), complicates the analysis of phenotype specificity. Here we generated an analytic pipeline that, while still indirect, provides specificity for a translation dependent outcome. We show that conditional depletion of *TgmS35* results in disruption of the activity of respiratory complex IV, which contains the mitochondrially encoded subunits (CoxI and CoxIII), at 48 hours after gene depletion (Figure 5A). On the other hand, complex V, for which the subunits are encoded in the nuclear genome, shows no decrease in activity at the 48-hour time point (Figure 5A). Complex V only shows a defect after a long period of *TgmS35* depletion, potentially due to secondary effects (Figure 5B). We further show that the assembly and activity of the TOM complex is unaffected at the 96-hour time point (Figure 5C). This approach of comparing the activities of complexes that do and do-not depend on mitochondrial translation for their assembly and activity provides a measure of mitochondrial translation in Apicomplexa and can be used in future studies to assess the involvement of other candidates with putative roles in mitochondrial translation, as well as the impact of drugs aimed at mitochondrial translation inhibition. Apicomplexan ribosomal proteins are not well studied. To our knowledge this is the first functional characterisation of a mitochondrial ribosomal protein in *T. gondii*. Our observations point to mitochondrial translational defect upon *TgmS35* depletion, thus supporting the role of *Tg*mS35 in the translation machinery. Finally, the data provides additional support to an active mitochondrial translation in these parasites.

In model organisms, *in vitro* functional assays performed with isolated mitochondria are extensively used to unravel mitochondrial functions including translation. Previously functional analysis of apicomplexan mitochondria has been hampered by the lack of good quality, large-scale, organelle enrichment protocols. This led to a reliance on whole cell biochemistry (e.g. Ke et al., 2018; Seidi et al., 2018) and on fluorescence microscopy analysis using fluorescent dyes (e.g. Vercesi et al., 1998; Garbuz and Arrizabalaga, 2017; Ovciarikova et al., 2017) for mitochondrial functional studies. Here, we developed an organelle enrichment protocol which generates high quantity fractions enriched with active mitochondria that can be used to investigate the activity of mitochondrial pathways *in vitro*. For example, enriched mitochondria are used for the complex V enzymatic activity performed using native-PAGE technique (Figure 5B and (Salunke et al., 2018)). The protocol we develop here for one of the most tractable apicomplexan parasites can be used to develop additional assays. By this, our work provides an avenue for prioritising mechanistic studies of these key conserved proteins. We found that the use of nitrogen cavitation to lyse *T. gondii* tachyzoites results in optimal lysis (Table S4), while maintaining any of the macromolecular-complexes examined here intact (TOM, complex IV, complex V and the putative mitochondrial small ribosome subunit (Figures 4B, 5B, C)). Nitrogen cavitation has previously been used for mitochondrial enrichment in *Plasmodium* (Mather et al., 2010; Kobayashi et al., 2007; Hata et al., 2018) and in the unrelated single cell eukaryote Trypanosomes (Schneider et al., 2007). We envisage this protocol will be invaluable for future efforts in the field to study *Toxoplasma* mitochondrial biochemistry, and to characterise other *Toxoplasma* organelles.

## Material and methods

### *In silico* mining

The mRNA abundance dataset generated by (Behnke et al., 2010) was queries via GeneSpring 11.0. Of the 14 homologs of mitochondrial protein import components (Table 1) 3 do not have cyclical expression pattern (Table S1.1). For each of the remaining 11, we searched for other genes with correlating cyclical expression patterns. We run Euclid and Spearman correlation tests at 0.9 and 0.95 scores and kept lists of correlation that were smaller than 70. This search identified 281 non-overlapping genes (the original search was performed with release 7.2 of ToxoDB, TGME49 strain; the same group of genes now correspond to 279 predicted genes from this strain)(search steps are shown in Table S1.2).

The orthology search leading to the 43 gene list was performed via the ToxoDB link to OrthoMCL. A search strategy queried for *Toxoplasma* genes with orthologs in *Plasmodium* 3D7 and with no orthologs in the three *Cryptosporidium* spp genomes available. The group of TGME49 genes that comply with these criteria was intersected with the 279 group resulting in 43 genes (Table S1.2, column C). Likewise, sorting based on fitness phenotypes scores was performed via the ToxoDB tool. All TGME49 genes were converted to their TGGT1 syntenic orthologs and then sorted according to their fitness scores.

The probability of export to the mitochondria was assessed using the MitoProt II algorithm (v.1.101) (Claros and Vincens., 1996) (https://ihg.gsf.de/ihg/mitoprot.html). Candidates with a MitoProt II score above 0.9 were considered to have a high probability of mitochondrial targeting, between 0.5-0.9 a medium probability, and below 0.5 a low probability. Export of mitochondria was also assessed using Predotar algorithm (v.1.04) (Small et al., 2004) (https://urgi.versailles.inra.fr/predotar/) with the “possibility of plastid targeting” option selected.

### Plasmid construct

For ligation independent cloning (LIC), between 500pb to 1.5kb homologous region of the 3’ end of each GOI was amplified by PCR from *T. gondii* RH genomic DNA with LIC sequence flanking these regions (primers are in Table S2). Products were LIC-cloned into linearised p3HA.LIC.CATΔpac containing a triple HA epitope tag as described previously (Huynh and Carruthers, 2009; Sheiner et al., 2011). The vectors were transfected into the TATi∆ku80 line (Sheiner et al., 2011) and selected with chloramphenicol.

For expression of tagged minigenes, the corresponding cDNAs were cloned (primers in Table S2) into the pDT7S4 expression vector that fuses a C-terminal Myc eptitope tag (van Dooren et al., 2008) via BglII and AvrII restriction sites, or into pTUB8mitoYFPFRB-DHFR vector that fuses a C-terminal YFP (Jimenez and Meissner unpublished, kind gift from Markus Meissner) via BstBI and NcoI restriction sites.

For promoter replacement, the ChopChoP (http://chopchop.cbu.uib.no/) tool was used to identify gRNAs covering the ATG of each GOI. The corresponding gRNAs were cloned into a U6 promoter and CAS9-GFP expressing vector (Tub-Cas9-YFP-pU6-ccdB-tracrRNA) (Curt et al., 2015) using the BsaI restriction site, and the final plasmid was purified by DNA midi-prep (Qiagen) per the manufacturer’s protocol. The DHFR selectable cassette and ATc repressible promoter were amplified by PCR from pDT7S4myc (van Dooren et al., 2008; Sheiner et al., 2011). Parasites were transfected with 50ug of the gRNA/CAS9 vector-PCR product mixture, and cassette integration was selected with pyrimethamine.

For transient gene disruption, each gene specific gRNA (sequences in Table S2) vector, synthesised by GenScript using backbone vector pU6-SAG1gRNA-DHFR (Serpeloni et al., 2016), was co-transfected with the CAS9-FLAG expressing vector (pU6-universal) (Sidik et al., 2016).

### Parasites culture

All *T. gondii* lines were cultured on Human Foreskin Fibroblast (HFFs) in DMEM media complemented with 10% FBS, 2% L-glutamine and 1% Penicillin/Streptomycin antibiotics and grown at 37°C with 5 % CO_2_ air atmosphere.

### Stable transfection

Linearised vectors were transfected into TATi∆ku80 (Sheiner et al., 2011) expressing line by electroporation. Parasites were selected by adding the appropriate drug to the culture medium for up to 2 weeks, then cloned on 96 well plates. Positive clones were screened by PCR using the appropriate primers listed in Table S2.

### Immunofluorescence assay and Microscopy

Parasites were inoculated onto human foreskin fibroblasts (HFFs) on coverslips for the mentioned time period and fixed with 4% paraformaldehyde (PFA) for 20 minutes at room temperature (RT). The PFA was washed away thrice with phosphate buffer saline (PBS). Cells were then permeabilised and blocked with blocking buffer (PBS, 0.2% Triton X-100, 2% Bovine Serum Albumin, BSA) for 20 minutes at RT. Cells were then labelled with different sets of primary and secondary antibodies: mouse anti-HA antibody (1:1000, Sigma), mouse anti-FLAG (1:1000), rabbit anti-TgMys (1:000) (Ovciarikova et al., 2017), mouse anti-Myc (1:1000, Cell Signalling), rabbit anti-TgTOM40 (1:2000)(van Dooren et al., 2016); coupled with goat anti-mouse or anti-rabbit fluorescent antibody (AlexaFluor 594 or 488 1:1,000 (Invitrogen)). Primary antibodies were incubated for 1 hour at RT. Coverslips were then washed thrice for 5 minutes with PBS-0.2% Triton X-100 and incubated with secondary antibodies for 45 minutes at RT in the dark. Coverslips were washed as above and mounted on microscope slides using DAPI-fluoromountG (Southern Biotech). Images were taken using a Delta Vision microscope as described (Ovciarikova et al., 2017).

### Mitochondrial morphology scoring

*CRISPR/Cas9 transient expression* - Parasites transiently co-expressing CAS9-FLAG expressing vector (pU6-universal) (Sidik et al., 2016) and gene specific sgRNA containing vector (Serpeloni et al., 2016), for genes *TGME49_214790*/*263680* and *226280*, were fixed 24 hours post-transfection. The CAS9-FLAG and mitochondria were visualised by immunofluorescence as described above. 50 vacuoles of CAS9-FLAG expressing parasites for each gene were then counted and mitochondrial morphology was assessed as followed: mitochondrial morphologies were sorted into two categories: (a) “normal” mitochondrial morphology (open, sperm-like morphologies (Ovciarikova et al; 2017)), (b) abnormal morphology (anything other than open and sperm-like morphologies), then scored. Dead parasites (displaying one or two balled mitochondria (Ovciarikova et al; 2017)) were not considered. Experiments were performed in triplicate.

*Conditional knockdown - rTgmS35*-3xHA mutant line was grown in the presence or absence of ATc 0.5 µM for 48 hours, fixed and mitochondria were visualized by immunofluorescence using anti-TgMys, as described above. Mitochondrial morphology was scored in the same manner as described for the above. Experiments were performed in triplicate, and a Chi-squared statistical test was performed.

### Plaque assay

1000, 300 and 100 parasites were inoculated in 6-well plates in duplicates, in the presence or absence of ATc 0.5 µM for 9 days. Cells were then fixed with 100% methanol for 5 minutes at RT, washed with PBS and crystal violet dye was added for 2 hours at RT to stain host cells, and then washed with PBS.

### Western Blot

Samples were resuspended in 1X NuPAGE LDS loading dye (Invitrogen) with 5% v/v beta-mercaptoethanol, then boiled at 95°C for 5 minutes and separated by SDS-PAGE. Proteins were transferred under wet conditions to nitrocellulose membrane (0.45 μm Protran™). Blots were labelled with the appropriate set of antibodies: primary rat anti-HA (1:500, Sigma), mouse anti-Myc (1:1000, Cell signalling) or rabbit anti-TgTOM40 (1:2000, (van Dooren et al., 2016)) antibodies coupled to secondary horseradish peroxidase (HRP)(Promega for mouse and rabbit, Abcam for rat) conjugated antibodies (1:10,000) or secondary fluorescent antibodies IRDye^®^ 800CW (1:10000, LIC-COR), and visualised using the Pierce ECL Western Blotting Substrate (Thermo Scientific) or the Odyssey LCX, respectively.

For the blue native-PAGE, proteins were transferred onto a PVDF membrane (0.45 μm, Hybond™) using wet transfer in Towbin buffer (0.025 M TRIS 0.192 M Glycine 10 % Methanol) for 60 minutes at 100 V. After transfer and immunolabelling revelation was carried out as described above with the Pierce ECL Western Blotting Substrate (Thermo Scientific).

### Protein import assay

Parasites were transfected with the Hsp60_L_-mDHFR-cMyc vector (van Dooren et al., 2016) (kind gift from Giel van Dooren), after growth in the presence or absence of 0.5 µM ATc for 24 and 72 hours. Transfected parasites were collected after an additional 24 hours growth in their respective treatment and western blot was performed using the respective parasite pellets as described above. Band intensity was measured using the ImageJ software and the ratio between the mature and pre-processed band intensity was calculated. A 1-way ANOVA statistical test was performed.

### Preparation of mitochondria enriched fraction

Parasites were cultured on HFFs in T150 flasks in the presence or absence of 0.5 µM ATc. Egressed parasites were collected (or, where necessary, parasites were scraped and syringed, e.g. at 96 hours) and filtered through a 3 µm pore size membrane. From this point on, all steps are carried out at 4°C or on ice. Parasites were centrifuged at 1500 × g for 15 minutes. Media was discarded and pellets were resuspended in PBS. Parasites in PBS were counted on a Neubauer chamber and centrifuged at 1500 × g for 15 minutes. The supernatant was discarded and the pellet resuspended in lysis buffer (50mM HEPES-KOH pH 7.4, 210 mM mannitol, 70 mM sucrose, 1 mM EGTA, 5 mM EDTA, 10 mM KCl, 1 mM DTT, 1 protease inhibitor cocktail tablet (Complete Mini, EDTA-free; Roche) per 50 ml) to a concentration of 5 × 10^8^ parasites mL^-1^. Parasites were transferred into a precooled nitrogen cavitation chamber and incubated at a pressure of 2750psi for 15 minutes on ice. After pressure release and centrifugation at 1500 × g for 15 minutes, the supernatant (lysate) was kept, and the pellet, containing unbroken parasites, was resuspended again at 5 × 10^8^ parasites mL^-1^ in lysis buffer. Repeating rounds of nitrogen cavitation were performed until >95% of parasite lysis was achieved (evaluated via counting using Neubauer chamber). Pooled lysates were span down at 1500 × g to remove any remaining unbroken parasites. The lysate was then aliquoted to the desired amount in microcentrifuge tubes and span down at 16,000 × g for 25 minutes. Supernatant discarded and pellet used immediately or stored at −80°C until use.

### Blue-Native Polyacrylamide Gel Electrophoresis

Whole parasite or previously aliquoted enriched mitochondria fraction were mixed with solubilisation buffer (750 mM aminocaproic acid, 50 mM Bis-Tris–HCl pH 7.0, 0.5 mM EDTA, 1 % (w/v) dodecyl maltoside) and incubated on ice for 5 minutes. The mixture was centrifuged at 16,000 × g at 4°C for 10 minutes. The supernatant containing solubilised membrane proteins was transferred into a new microcentrifuge tube and 0.25% (w/v) Coomassie G250 (final concentration) was added. The anode buffer (50 mM Bis-Tris–HCl pH 7.0) and cathode buffer (50 mM Tricine, 15 mM Bis-Tris–HCl pH 7.0, 0.02% Coomassie G250 (Serva)) were poured into their respective tank compartment and the appropriate amount of protein (55μg or 5×10^5^ parasites) was loaded per lane on a NativePAGE™ 4-16% Bis-Tris Gel (Novex-Life technologies). 5 μl NativeMark™ (Invitrogen) was used as a molecular weight marker. Gels were run for ~45 minutes at 100 V, 4-10 mA at 4°C with cathode buffer containing 0.02% (w/v) Coomassie G250, then for ~2.5 hours at 250 V, 15 mA with cathode buffer containing 0.002% (w/v) Coomassie G250, until the dye front reached the bottom of the gel.

### High resolution Clear-Native Polyacrylamide Gel Electrophoresis

Carried out as described in (Wittig et al., 2007) with minor modifications. Briefly, Whole parasite or previously aliquoted mitochondria enriched fraction (75 µg) were mixed with solubilisation buffer (50 mM NaCl, 50 mM Imidazole, 2 mM 6-aminohexanoic acid, 1 mM EDTA – HCl pH 7.0, 2% (w/v) n-dodecylmaltoside) and incubated on ice for 10 minutes. The mixture was centrifuged at 16,000 × g at 4°C for 15 minutes. A final concentration of 6.25% glycerol and 0.125% Ponceau S was added to the solubilised membrane proteins. The anode buffer is composed of 25 mM Imidazole – HCl pH to 7.0 and the cathode buffer of 50 mM Tricine, 7.5 mM Imidazole, 0.02% w/v n-dodecylmaltoside, 0.05% sodium deoxycholate. Gels were run for 30 minutes at 100 V, 10 mA, then 250-300 V, 15 mA at 4°C until it the dye front reached the bottom.

### In-gel activity staining

Activity stains were carried out as described previously (Sabar et al., 2005). Briefly, gels were equilibrated in buffer without staining reagents for 10 minutes. For complex IV, oxidation activity was shown using 50 mM KH_2_PO_4_, pH 7.2, 1 mg mL^-1^ cytochrome *c*, 0.1% (w/v) 3,3’-diaminobenzidine tetrahydrochloride. Stains were visible after 30 minutes. Staining was continued with pictures taken of the stained gels at regular intervals. For complex V, ATP hydrolysis activity was visualised using 35 mM Tris, 270 mM glycine, pH 8.4, 14 mM MgCl_2_, 11 mM ATP, 0.3% (w/v) Pb(NO_3_)_2_. Stains were visible after 1 hour. Pictures were taken against a black background for optimal visualisation of white lead precipitates.

### Mass spectrometry

Proteins were identified using nanoflow HPLC electrospray tandem mass spectrometry (nLC-ESI.MS/MS) at Glasgow Polyomics. Tryptic peptides, generated using in-gel digest procedure (Williams et al., 2018), were analysed as previously described (Akpunarlieva et al., 2017). Protein identities were assigned using the Mascot search engine (v2.6.2, Matrix Science) to interrogate protein sequences in the *T. gondii* genome sequence dataset, ToxoDB 35 release. During result analysis, only peptides with Mascot score of 20 and above (namely the probability that this match might be a random event is 10^-2^ or lower) were included in the analysis.

## Acknowledgements

We thank Glasgow Polyomics, and Dr. Richard Burchmore for advice on sample preparation and MS data analysis. We thank Markus Meissner for sharing reagents and for productive discussions and advice. We thank Giel van Dooren and Mohamed-Ali Hakimi for sharing plasmids. There are no conflicting interests. LS is a Royal Society of Edinburgh Personal Research Fellow. The work is funded by BBSRC BB/N003675/1 and by Wellcome Trust Institutional Strategic Support Fund Fellowship to LS.

## Author contributions

L.S. conceived the research; L.S. A.L. and A.M developed the method; A.L. A.M J.O. J.T. and L.S. performed the experiments; L.S. A.L. and A.M. discussed the data and wrote the paper.

