## supplementary figures 1 and 2 for "*In silico* screen identifies a new *Toxoplasma gondii* mitochondrial ribosomal protein essential for mitochondrial translation"

**Supplementary figures and tables:**

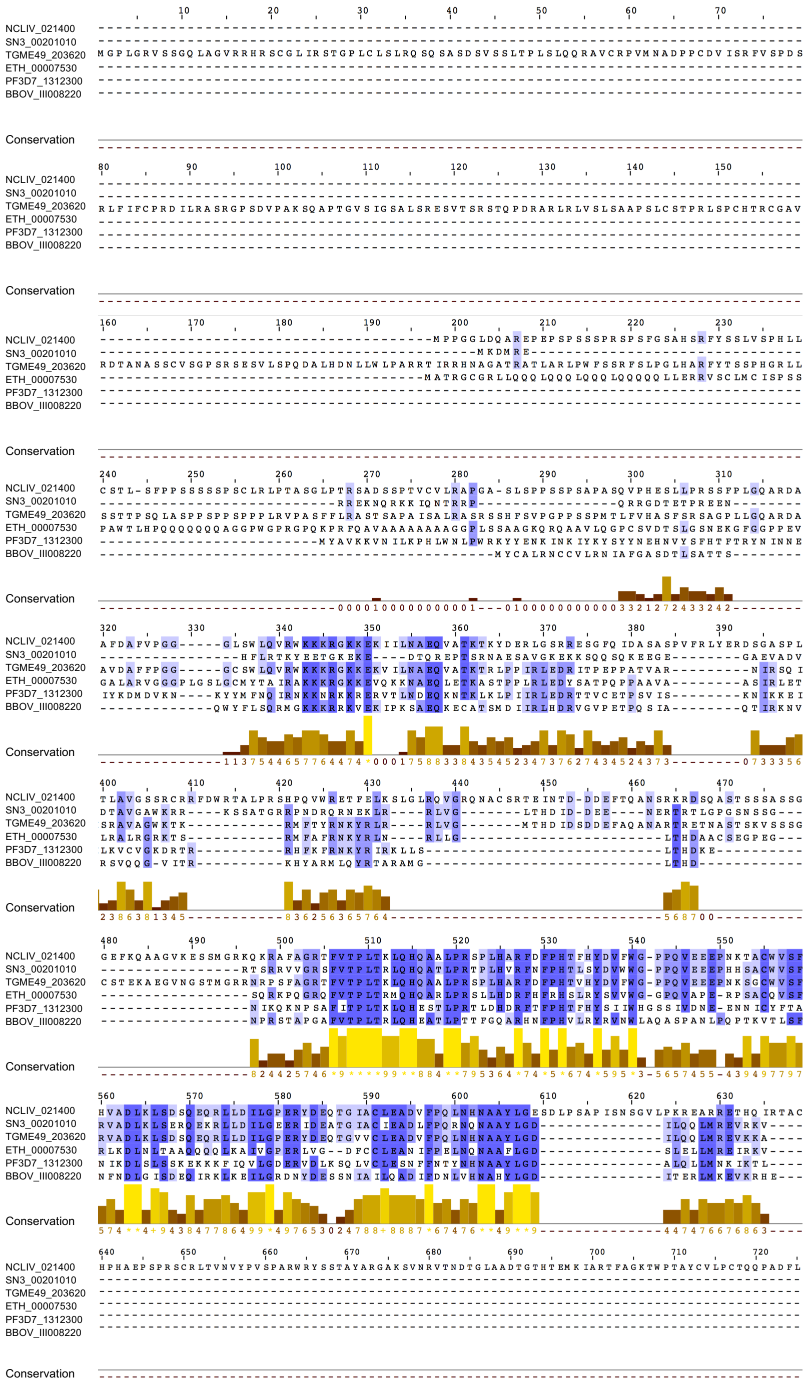


**Supplemental Figure 1. Protein alignment of *Tg*mS35 apicomplexan homologs.** Homologs identified via BLAST in EUpathDB.org. Alignment performed using clustalX pairwise alignment and consensus colours generated via Jalview. Amino acid conservation in shown within the alignment in a blue-scale whereby light is low similarity and dark blue is high similarity. Conservation is also shown as bars at the bottom. Bar colour is yellow-scale, whereby light yellow is high similarity and dark orange/brown is low conservation.

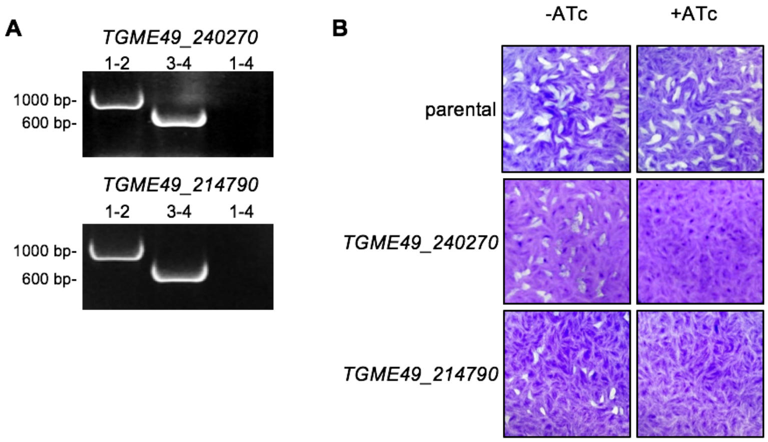


**Supplemental Figure 2. Down-regulation of genes *TGME49_240270* and *TGME49_214790* results in parasite growth defect. (A)** Validation of the promoter integration in each gene locus via PCR analysis using primers 1, 2, 3, and 4 represented in the scheme in figure 4C **(B)** Plaque assays performed in the absence (-) or presence (+) of ATc after 9 days. Parasite line name is indicated on the left. The image of parental line plaque assay is the same image of parental line used in figure 4G.
